# Engineered Wnt7a ligands rescue blood brain barrier and neurobehavioral deficits in a mouse model of COVID-19

**DOI:** 10.1101/2022.06.02.494552

**Authors:** Troy N. Trevino, Avital B. Fogel, Jacob Class, Mark A. Sanborn, Benoit Vanhollebeke, Jalees Rehman, Leon M. Tai, Justin M. Richner, Sarah E. Lutz

**Affiliations:** Department of Anatomy and Cell Biology, University of Illinois at Chicago, College of Medicine, Chicago, IL 60612, USA; Department of Microbiology and Immunology, University of Illinois at Chicago, College of Medicine, Chicago, IL 60612, USA; Department of Biochemistry and Molecular Genetics, University of Illinois at Chicago, College of Medicine, Chicago, IL 60612, USA; Laboratory of Neurovascular Signaling, Department of Molecular Biology, ULB Neuroscience Institute, Université libre de Bruxelles (ULB), Gosselies B-6041, Belgium

**Author notes:** Corresponding author: Sarah E. Lutz, PhD. Department of Anatomy and Cell Biology, University of Illinois at Chicago College of Medicine, 808 S. Wood St., Rm 578 MC 512, Chicago IL 60612., **Email:**. **Author Contributions:** TNT, BV, JR, JMR, SEL designed research. TNT, ABF, JC, MAS, JMR, SEL performed research. TNT, ABF, JC, MAS, LMT, SEL analyzed data. TNT, BV, JR, LMT, JMR, SEL wrote the paper. **Competing Interest Statement:** SEL and BV filed a US provisional application on compositions for preventing and/or treating NeuroCOVID. BV is a founder and shareholder of NeuVasQ Biotechnologies.

**Keywords:** Blood-brain barrier, SARS-CoV-2, Wnt7a, endothelial cell, neuroinflammation

## Abstract

Respiratory infection with SARS-CoV-2 causes systemic vascular inflammation and cognitive impairment. We sought to identify the underlying mechanisms mediating vascular dysfunction and inflammation following mild respiratory SARS-CoV-2 infection. To this end, we conduced unbiased transcriptional analysis to identify brain endothelial cell signaling pathways dysregulated by SARS-CoV-2 *in vivo*. This analysis revealed significant suppression of Wnt/β-catenin signaling, a critical regulator of blood brain barrier integrity. We therefore hypothesized that enhancing cerebrovascular Wnt/β-catenin activity would offer protection against BBB permeability, neuroinflammation, and neurological signs in acute infection. Indeed, we found that delivery of cerebrovascular-targeted, engineered Wnt7a ligands protected blood brain barrier integrity, reduced T cell infiltration of the brain, and reduced microglial activation in SARS-CoV-2 infection. Importantly, this therapeutic strategy also mitigated SARS-CoV-2 induced deficits in the novel object recognition assay for learning and memory and the pole descent task for bradykinesia. These observations suggest that enhancement of Wnt/β-catenin signaling or its downstream effectors could be potential interventional strategies for restoring cognitive health following acute viral infections.

## Introduction

Respiratory viral infections are increasingly recognized to influence neurological function(1). This has been highlighted in the COVID-19 pandemic, which has resulted in widespread acute and chronic neurologic deficits. A prominent feature of acute SARS-CoV-2 infection is endothelial inflammation (1). There is increasing evidence that cerebrovascular endothelial cells can also become inflamed and dysfunctional in COVID-19 (1). One major function of brain endothelial cells is to regulate the permeability of the cerebrovasculature to macromolecules and leukocytes, a feature known as the blood-brain barrier (BBB). BBB disruption contributes to neuroinflammation and cognitive impairment. Many groups have reported evidence of BBB disruption in COVID-19 patients and in animal models of SARS-CoV-2 infection (2-5). However, the extent to which brain endothelial dysfunction contributes to neurological deficits, and the potential pathways underlying this, in acute SARS-CoV-2 have not yet been fully defined. Using an unbiased transcriptomic approach, we determined that respiratory SARS-CoV-2 infection dysregulated brain endothelial Wnt/β-catenin signaling, a critical regulator of BBB function. Targeted delivery of an engineered Wnt7a ligand to the brain vasculature prevented BBB disruption and restored neurobehavioral function in SARS-CoV-2 infection.

## Results

We intranasally inoculated C57Bl/6 mice with mouse-adapted SARS-CoV-2 strain MA10 or vehicle and euthanized at 4 days post inoculation (DPI). Twelve-month old C57Bl/6 mice infected with MA10 develop a mild respiratory illness and recapitulate features of cerebrovascular dysfunction and neuroinflammation observed in patients (6, 7). We conducted RNA-seq on brain endothelial cells of mice with and without SARS CoV-2 infection (Figure 1A-C). SARS-CoV-2 infection upregulated gene expression pathways related to response to virus, cell substrate adhesion, and apoptosis, and downregulated pathways related to extracellular matrix organization and immunosuppression. Importantly, we found that SARS-CoV-2 infection downregulated genes in the Wnt signaling pathway in brain endothelial cells (Figure 1D). This finding was important because the Wnt/β-catenin signaling pathway is required for blood-brain barrier (BBB) integrity (8-12). Canonical Wnt ligands causes a signaling cascade resulting in β-catenin dependent transcription of target genes (8, 9). Further analysis in the Kyoto Encyclopedia of Genes and Genomes (KEGG) Wnt/β-catenin signaling pathway (Figure 1E) highlighted that downstream targets of Wnt/β-catenin signaling were also downregulated by respiratory SARS-CoV-2. These included the transcriptional target Ccn4/WISP1, transcriptional coactivators and effectors Sox9 and Foxo1, and pathway regulator Dkk2 (Figure 1E). Of all targets in the KEGG Wnt/β-catenin signaling pathway, 17 genes were significantly downregulated and 4 were significantly upregulated by SARS-CoV-2 (Figure 1E). As a validation of changes in the Wnt/β-catenin signaling pathway, we assessed a downstream transcriptional target of canonical β-catenin transcription in brain endothelial cells, Nkd1 at the protein level and found concordant downregulation (Figure 1F-H). Together, these data suggest that the Wnt/β-catenin signaling pathway is dysregulated in brain endothelial cells following SARS-CoV-2 infection.

**Figure 1.**
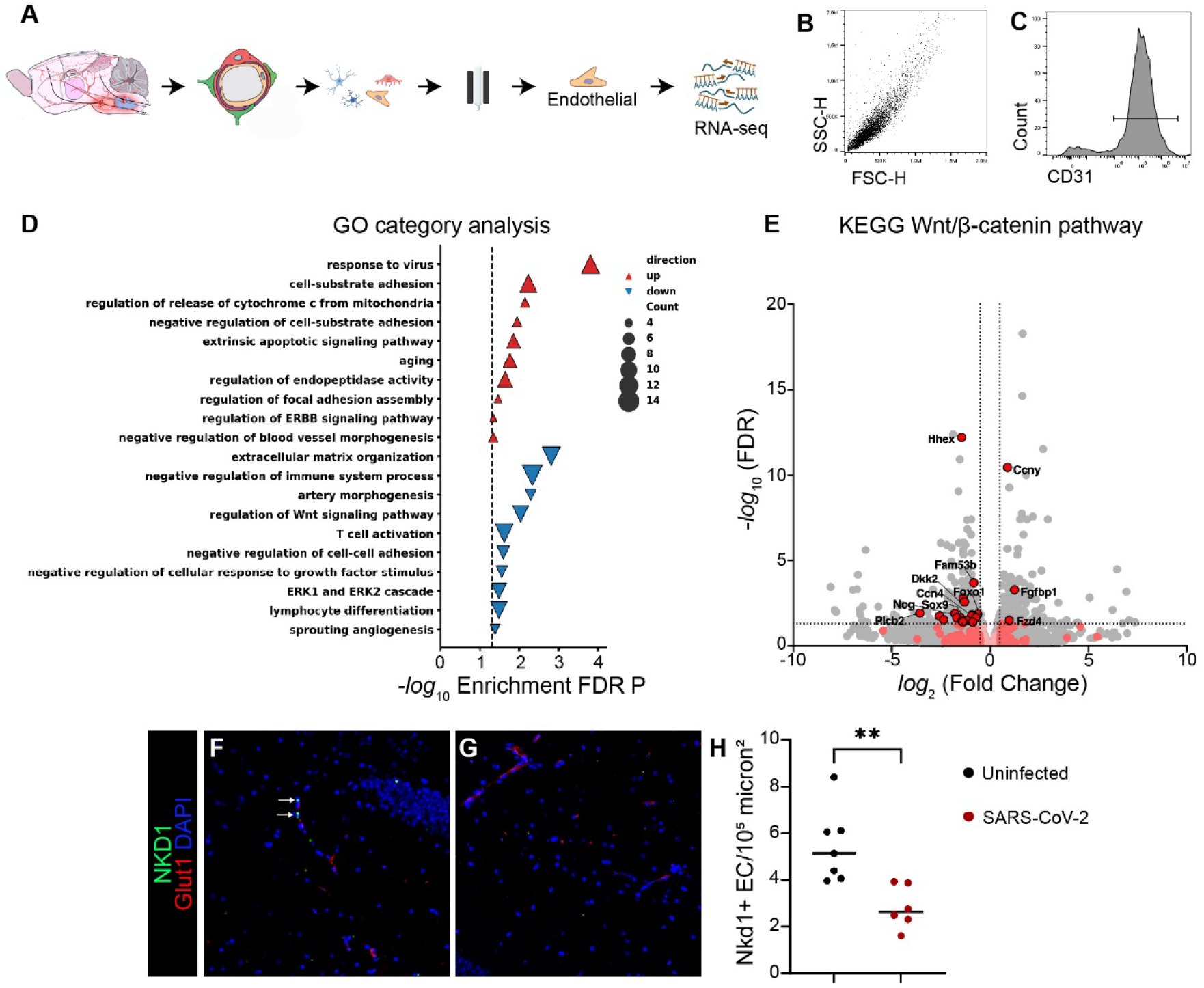
Dysregulation of brain endothelial cell Wnt/β-catenin signaling in SARS-CoV-2 infection. A) Four days after respiratory inoculation with SARS-CoV-2, mice were euthanized, brains were dissected, microvessel fragments isolated, and negative and positive column-based selection were deployed to isolate endothelial cells for bulk RNA-sequencing. B) Flow cytometric analysis of isolated brain endothelial cells demonstrating uniform population for side-scatter (SSC-H) and forward scatter (FSC-H). C) Flow cytometric analysis showing histogram for CD31 out of all cells in panel B. 94% of cells were positive for the endothelial cell marker CD31. D) Gene ontology category analysis of differentially expressed genes in brain endothelial cells of mice with SARS-CoV-2 as compared to mock infection. E) Volcano plot in which differentially expressed genes in KEGG Wnt/β-catenin pathway are depicted in red. Non-differentially expressed KEGG Wnt/β-catenin genes are depicted in pink. Gray dots represent transcripts that are not part of the KEGG Wnt/β-catenin pathway. F-G) Immunostaining for the β-catenin transcriptional target Nkd1 (green) in brain sections of SARS-CoV-2 or mock-infected mice. Brain endothelial cells are visualized with Glut-1 (red). Nuclei are visualized with DAPI (blue). Monochromatic images are provided as Supplemental Data. H) Quantification of density of Nkd1+ endothelial cells in the hippocampus of SARS-CoV-2 or mock-infected mice. N=4-5 mice/group. Unpaired student’s t-test, \*\**p*<0.01.

We next explored the extent to which enhancing brain endothelial Wnt/β-catenin pathway activity could improve BBB function, neuroinflammation, and neurological signs SARS-CoV-2 infection. To do this we chose an agonist-based approach. Our rationale was that because Wnt7a receptors Gpr124, Reck, Fzd4, and Lrp4/5 (8, 9) were not among the pathway genes downregulated by SARS-CoV-2 infection (Figure 1E), brain endothelial cells might retain responsiveness to receptor agonism. We deployed the cerebrovascular-tropic gene therapy vector AAV:PHP.eB to deliver Wnt7a engineered with a lysine to alanine mutation at position 190 (Wnt7a^K190A^) to specifically activate Gpr124/Reck (8). Gpr124/Reck is the receptor used by endothelial cells to signal downstream of Wnt7a; therefore Wnt7a^K190A^ has specificity for brain endothelial cells. We previously demonstrated that cerebrovascular-targeted AAV-PHP.eB-Wnt7a^K190A^ did not impact behavior or BBB in healthy adult mice, and had therapeutic efficacy in mouse models of glioblastoma and ischemic stroke by preventing BBB leakage (8). We used AAV:PHP.eB encoding green fluorescent protein (GFP) as a vehicle control. We administered AAV-PHP.eB-Wnt7a^K190A^ or AAV:PHP.eB-GFP to mice 18 days prior to inoculation with SARS-CoV-2. Consistent with our previous studies, we found that ∼25% of microvessel-associated cells were transduced, as assessed by flow cytometry for GFP (Figure 2A-B). Weight loss was not significantly different between groups (Figure 2C).

**Figure 2.**
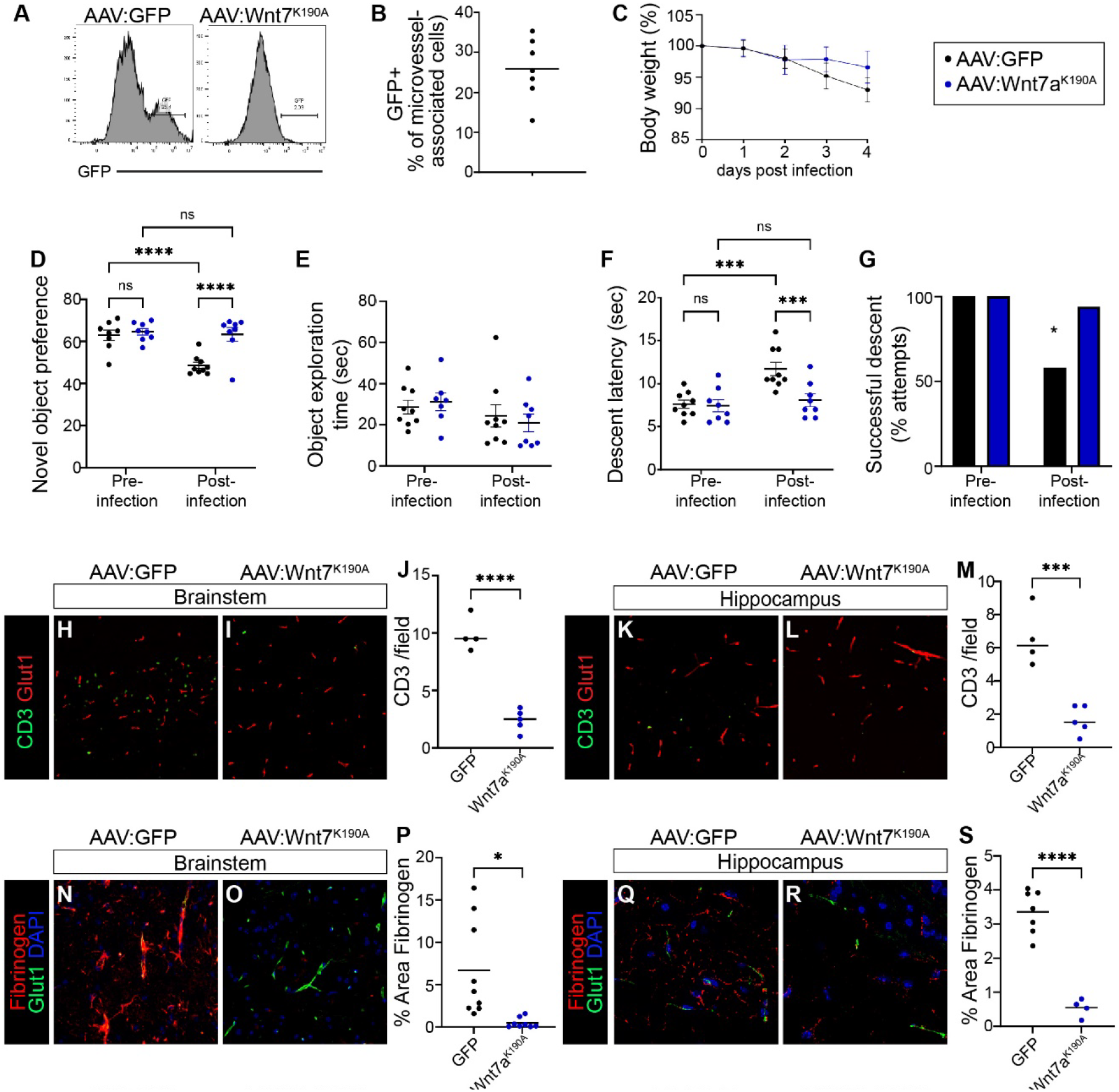
Cerebrovascular-targeted engineered Wnt7aK190A protects against neurobehavioral impairment, neuroinflammation, and BBB leakage in SARS-CoV-2 infection. A) Flow cytometric histograms of GFP fluorescence in microvessel-enriched fragments of brains of mice treated with AAV-PHP.eB-GFP or AAV-PHP.eB-Wnt7a^K190A^. B) 25% of microvascular cells are positive for GFP in samples from brain of mice treated with AAV-PHP.eB-GFP. 8-9 mice/group. C) Body weight changes after SARS-CoV-2 infection did not significantly differ between mice receiving AAV-PHP.eB-Wnt7a^K190A^ and mice receiving control vector (area under curve, *p* >0.05). 8-9 mice/group. D) Mice were treated with AAV-PHP.eB-GFP or AAV-PHP.eB-Wnt7a^K190A^ 18 days prior to inoculation with SARS-CoV-2. The novel object recognition test for learning and memory was conducted 2 days prior and 5 days after SARS-CoV-2 infection. There was a Treatment [F_(1, 29)_ = 13.05, *p* = 0.0011], SARS-CoV-2 [F_(1, 29)_ = 11.75, *p* = 0.0018], and Treatment x SARS-CoV-2 interaction [F_(1, 29)_ = 8.397, *p* = 0.0071] for novel object preference. Post hoc analysis revealed that SARS-CoV-2 infection impaired NOR in mice treated with the control GFP vector (*p* <0.0001) but not in mice treated with Wnt7a^K190A^ (*p* = 0.7145). Indeed, SARS-CoV-2 infected mice treated with Wnt7a^K190A^ performed significantly better than SARS-CoV-2 infected mice treated with the control vector (*p* <0.0001). E) Total object exploration was not significantly influenced by Treatment [F_(1, 29)_ = 0.009622, *p* = 0.9225], SARS-CoV-2 [F_(1, 29)_ = 2.679, *p* = 0.1125], or Treatment x SARS-CoV-2 interaction [F_(1, 29)_ = 0.4577, *p* = 0.5041], indicating that the novel object recognition task was not confounded by inactivity. F) Pole descent latency, a measure of motor coordination impairment, was significantly influenced by Treatment [F_(1, 30)_ = 7.855, *p* = 0.0088], SARS-CoV-2 [F_(1, 30)_ = 12.10, *p* = 0.0016], and Treatment x SARS-CoV-2 interaction [F_(1, 30)_ = 6.410, *p* = 0.0168]. Post-hoc analysis revealed the difference was driven by impairment in the vector-treated SARS-CoV-2 group as compared to the Wnt7a^K190A^-treated group (*p* = 0.0007 or the uninfected group (*p* = 0.0001). G) SARS-CoV-2 infected mice treated with AAV-PHP.eB-Wnt7a^K190A^ fell off the pole significantly fewer times than those treated with AAV-PHP.eB-GFP (Fisher’s exact test, *p* = 0.0198). H-M) Immunostaining for CD3+ T cells (green) in brainstem (H, I) and hippocampus (K, L) of SARS-CoV-2 infected mice treated with AAV-PHP.eB-Wnt7a^K190A^ or AAV-PHP.eB-GFP. Tissue was collected 5 days after SARS-CoV-2 infection. Glut1 (red) was used to visualize endothelial cells. SARS-CoV-2 infected AAV-PHP.eB-Wnt7a^K190A^ treated mice had significantly lower density of CD3+ T cells than did AAV-PHP.eB-GFP treated mice in brainstem (J) and in hippocampus (M). N=4-5 mice/group, unpaired Student’s t-test. N-S) Immunostaining for fibrinogen (red) in brainstem (N, O) and hippocampus (Q, R) of SARS-CoV-2 infected mice treated with AAV-PHP.eB-Wnt7a^K190A^ or AAV-PHP.eB-GFP. Glut1 (green) was used to visualize endothelial cells. SARS-CoV-2 infected AAV-PHP.eB-Wnt7a^K190A^ treated mice had significantly lower perivascular fibrinogen than did AAV-PHP.eB-GFP treated mice in brainstem (P, unpaired Student’s t-test) and in hippocampus (S, unpaired Student’s t-test with Welch’s correction for unequal variances). N=4-9 mice/group. \**p*<0.05, \*\**p*<0.01, \*\*\**p*<0.001, \*\*\*\**p*<0.0001. Monochromatic images are provided as Supplemental Data.

A complication of acute SARS infection is neurological impairment (1). Therefore, we initially evaluated whether cerebrovascular-targeted Wnt7a^K190A^ might offer protection against neurological signs induced by SARS-CoV-2 in mice. We conducted the novel object recognition test, an assay of learning and memory related to hippocampal function, in mice with AAV-PHP.eB-Wnt7a^K190A^ or AAV:PHP.eB-GFP 2 days before SARS-CoV-2 infection and 5 days after SARS-CoV-2 infection. Acute SARS-CoV-2 caused significant impairment in the novel object recognition test (Figure 2D). Importantly, AAV-PHP.eB-Wnt7a^K190A^ mitigated SARS-CoV-2 induced deficits in the novel object recognition task (Figure 2D). Total exploration was not significantly affected by AAV nor by SARS-CoV-2 (Figure 2E), indicating that the decrease in novel object preference was not driven by potentially confounding sickness behaviors or motor deficits. Also as expected, no preference was noted in the training phase (object preference score 50.5+/-7.0%, n=16). Some COVID-19 patients experience autonomic dysfunction. We assessed this in mice with the pole descent task, which requires brainstem integration of proprioceptive sensation and cortical input. SARS-CoV-2 significantly prolonged pole descent time (Figure 2F) and increased the incidence of falls (Figure 2G). Importantly, AAV-PHP.eB-Wnt7a^K190A^ mitigated SARS-CoV-2 induced deficits in the pole descent task (Figure 2F-G). These results indicate that cerebrovascular-targeted engineered Wnt7a^K190A^ significantly improved neurological outcome of SARS-CoV-2 infection.

BBB dysfunction can lead to neuroinflammation and neurological problems. We next examined whether AAV-PHP.eB-Wnt7a^K190A^ prevented BBB dysfunction in SARS-CoV-2 infection. Two markers of BBB dysfunction are leukocyte infiltration and high levels of blood proteins in the brain. We therefore measured CD3+ T cells and fibrinogen extravasation in hippocampus and brainstem. We observed ∼80% fewer CD3+ T cells in the brain parenchyma in hippocampus and brainstem in mice that received AAV-PHP.eB-Wnt7a^K190A^ prior to SARS-CoV-2 as compared to those receiving vector control prior to SARS-CoV-2 (Figure 2H-M). We also observed ∼80% less perivascular fibrinogen in the hippocampus and brainstem in mice that received AAV-PHP.eB-Wnt7a^K190A^ prior to SARS-CoV-2 as compared to those receiving vector control prior to SARS-CoV-2 (Figure 2N-S). These data indicate that cerebrovascular-targeted canonical Wnt activation prevents BBB leakage and neuroinflammation caused by SARS-CoV-2.

## Discussion

SARS-CoV-2, like other respiratory infections, can cause substantial neurological impairment (1, 13). However, because such infections cause systemic inflammation, it has been difficult to determine the specific contribution of brain endothelial dysfunction. BBB leakage, leukocyte infiltration, microglial activation, and neuronophagia are well documented in COVID-19 clinical cases and in animal models (1-3, 5, 7, 13, 14), and are plausible causes of neurological signs of disease. Our study provides direct evidence that modulating brain endothelial cell function can mitigate neuroinflammation and neurobehavioral impairment caused by acute SARS-CoV-2 infection.

In this study, our goal was to use an unbiased transcriptomic approach to identify brain endothelial cell gene expression pathways that are dysregulated following respiratory SARS-CoV-2 infection and could be therapeutically targeted. We identified dysregulation of Wnt/β-catenin signaling, which maintains BBB integrity by promoting tight junctions, glucose transporters, pericyte/ECM coverage, and suppressing caveolar transport (8-12). Brain endothelial cell β-catenin signaling is neuroprotective in mouse models for multiple sclerosis by suppressing T cell infiltration, demyelination, and mortality (12). Furthermore, cerebrovascular-targeted Wnt ligands engineered for stable endothelial activation protect against glioblastoma and ischemic stroke (8). Supported by these findings, we used a gene therapy approach to target brain endothelial cell Wnt/β-catenin signaling in a mouse model for acute COVID-19. We determined that cerebrovascular-targeted delivery of Wnt7a receptor agonists with enhanced specificity brain endothelial cells prevented learning and memory deficits, bradykinesia, and BBB leakage in mice infected with SARS-CoV-2. Future studies are warranted to understand the proximal mechanisms by which endothelial β-catenin signaling reduces neurobehavioral deficits in SARS-CoV-2, and to interrogate whether restoring Wnt/β-catenin signaling can also prevent long-term neurological complications of respiratory SARS-CoV-2.

## Materials and Methods

### Mice

Twelve-month-old C57Bl/6 mice (Jackson laboratories) on standard light-dark cycles with ad libitum food and water were transferred to Animal BioSafety Level 3 facilities and micro-isolation cages (3-5 mice/cage) >2 days prior to inoculation and randomized to intranasal inoculation with 1 × 10^4^ foci-forming units (FFU) SARS-CoV-2 mouse adapted MA10 in 25 μl PBS or vehicle (saline). Animal studies were approved by the Animal Care Committee at UIC (20-107; 20-160). MA10 was provided by Ralph Baric (University of North Carolina, Chapel Hill, North Carolina, USA) (6). SARS-CoV-2 (MA10) was propagated and titered on Vero-E6 cells expressing ACE2 and TMPRSS2 (ATCC, CRL1586).

### Modulation of Wnt signaling

AAV-PHP.eB-Wnt7a^K190A^P2A-GFP and AAV-PHP.eB-GFP (8) were administered to mice by retroorbital injection of 2x10^11^ viral genomes in 25 μl of PBS 18 days prior to inoculation with SARS-CoV-2 MA10 (8). Behavioral testing and euthanasia was conducted at 5 DPI.

### Endothelial cell isolation

Mice were transcardially perfused with PBS. Brainstem microvascular endothelial cells were isolated using gradient centrifugation (15). Microvessels were dissociated with collagenase/dispase (Millipore Sigma 10269638001), DNase (Worthington LK003172) for 1 h in 37ºC water bath and passed through 100 μm cell strainer (PluriSelect USA 43-10100-60). Dissociated microvascular cells were stained for flow cytometric analysis or were additionally processed with myelin removal beads (Miltenyi 130-069-731) and 3 sequential positive selections with CD31 microbeads (Miltenyi 130-097-418) for RNA isolation (Qiagen RNeasy Micro Kit 74004). cDNA library preparation used Oligo-dT at 60 million clusters/sample (University of Chicago Genomics Facility).

### RNA-sequencing and analysis

Fastq files were quality checked using FastQC (https://www.bioinformatics.babraham.ac.uk/projects/fastqc/)prior to downstream analysis. Reads were aligned using STAR (version 2.7.6a) against the GRCm38 (mm10) genome provided by Ensembl. Count tables were generated using featureCounts (Subread release 2.0.1). Differential expression analysis was performed between testing groups using Deseq2. Clusterprofiler was used for overrepresentation analysis of the differentially expressed genes against the gene ontology database. RNAsequencing data has been deposited on GEO.

### Flow cytometry

Cells were sequentially incubated with viability dye (Zombie Green, Biolegend 423111), Fc receptor blockade (Biolegend 101319), and anti-CD31 antibody (BD Biosciences 561073). Cells were fixed (Biolegend 421403) and analyzed (Beckman CytoFLEX S).

### Behavioral Assays

Behavior tasks were conducted in a dark biosafety cabinet laminar flow hood in the BSL3 facility. Novel object recognition test was conducted between 7:00-10:00AM by filming mice for ten minutes with an overhead camera (Logitech C920S HD) in white plastic bins 13 inches x 19 inches (Ikea) with pebbled floor at 5DPI. Intersession interval was 12h. Preference was (sec investigating novel object)/(sec investigating any object)*100. 50% indicates no preference. For the pole descent assay, the rod of a buret support stand (½ inch diameter, 18 inches length) was mounted on a metal base covered with corn cob bedding. Average latency to descend was measured in two consecutive trials. Falls were scored as failure to descend and excluded from latency averages.

### Histology

Mice were perfused with a peristaltic pump. Brains were fixed in 4% paraformaldehyde and paraffin embedded. Antigen retrieval (10 mM Tri-sodium citrate [dihydrate], 0.05% Tween-20, pH 6.0; or Tris-EDTA pH 9.0 for Iba1) was 40 minutes at 98°C. Slides were blocked with 5% normal goat serum, 0.1% Triton-X 100. Primary antibodies incubated overnight at 4°C at 1:100 included CD3 (Cell Signaling 78588), Glut1 (Abcam ab40084), Fibrinogen (LS Bio LS-C150799-1), Nkd1 (Bioss 19005), CD68 (Biolegend FA-11), and Iba1 (Abcam ab178847). Alexa-fluorophore conjugated secondary antibodies (Invitrogen) were incubated 2h at 22°C. Mounting medium contained DAPI (Ibidi 50011). Microscopy was conducted using Zeiss LSM880 or Leica DMI8 microscopes. Quantification was performed using FIJI (NIH).

### Statistics

Statistics were conducted using GraphPad Prism 10. Pairwise comparisons used Student’s t-test, with Welch’s correction when variances were unequal. % Body weight was assessed by area under the curve. Pole descent failure was calculated with two-sided Fisher’s exact (Chi-square) test. Other behavioral tasks were tested with two-way ANOVA; significant interactions were compared by Sidak’s multiple comparisons test.

## Acknowledgments

This work was supported by the DOD MS200290 and NIH KL2TR002002 (SEL), T32HL139439 (TNT and MAS), R01AG061114 (LMT), R61NS114353 (LMT), R01AI150672 (JMR), R01HL162308 (JR), the ERC to BV (Ctrl-BBB 865176), and University of Illinois Institutional Funds (SEL and LMT). Research support services were obtained from the Research Histology Core and the Center for Clinical and Translational Science Biostatistics Core at the University of Illinois at Chicago. SEL and BV filed a US provisional application on compositions for preventing and/or treating neurocovid. BV is a founder and shareholder of NeuVasQ Biotechnologies.

## Figures

**Supplemental Figure 1:**
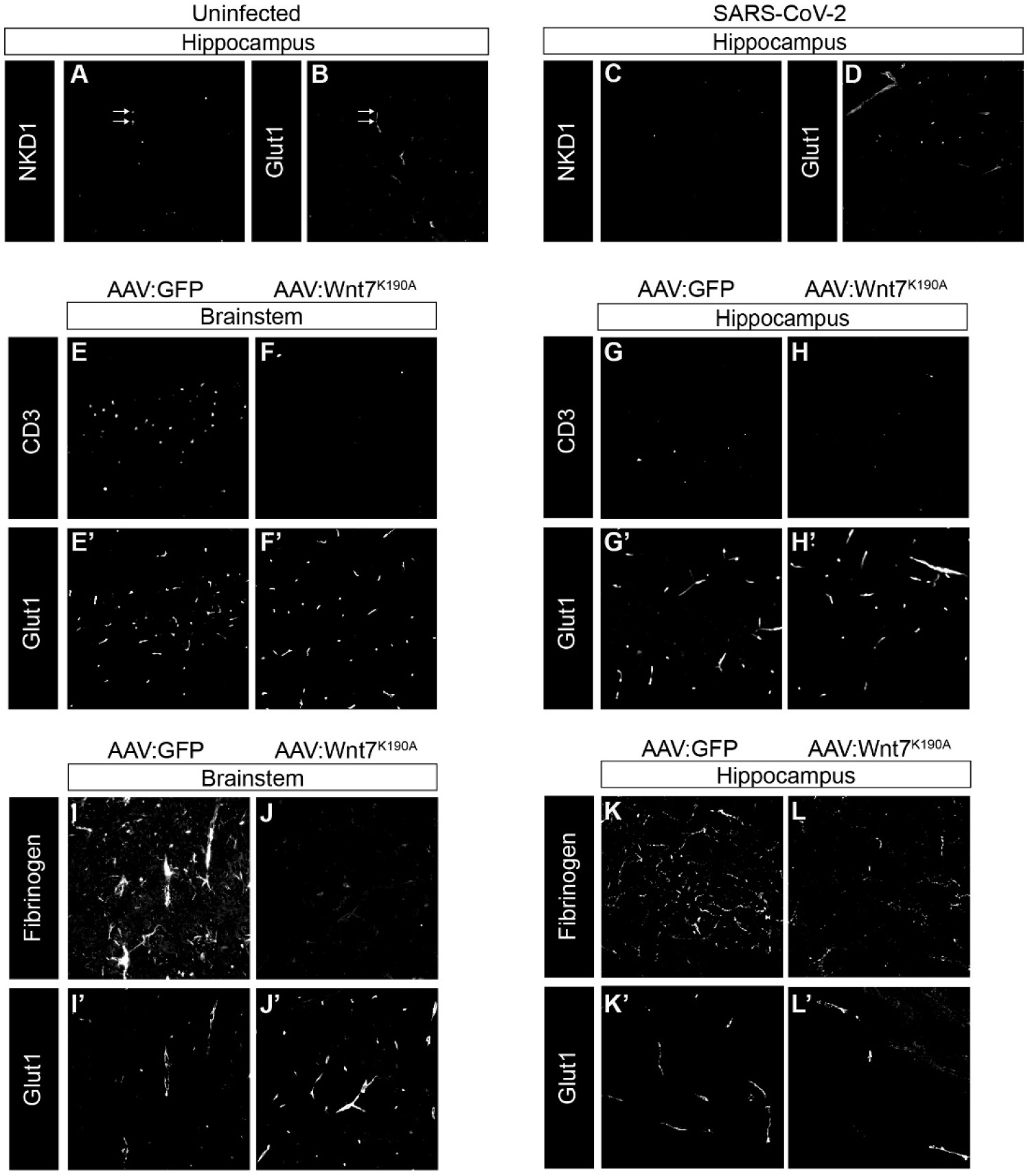
Monochromatic representations of micrographs shown in the main figures. A-D) Immunostaining for the β-catenin transcriptional target Nkd1 (A, C) in hippocampus sections of uinfected (A-B) or SARS-CoV-2-infected (C-D) mice. Brain endothelial cells are visualized with Glut-1 (B, D). E-H) Immunostaining for CD3+ T cells in brainstem (E, F) and hippocampus (G, H) of SARS-CoV-2 infected mice treated with AAV:GFP (E, G) or AAV:Wnt7a^K190A^ (G, H). Brain endothelial cells are visualized with Glut-1 (E’-H’). I-L) Immunostaining for the blood protein fibrin (fibrinogen) in brainstem (I, J) and hippocampus (K, L) of SARS-CoV-2 infected mice treated with AAV:GFP (I, K) or AAV: Wnt7a^K190A^ (J, L). Brain endothelial cells are visualized with Glut-1 (I’-L’).

## Notes

### Summary of Updates

We have updated the manuscript with (1) RNAsequencing data indicating that the Wnt/beta-catenin signaling pathway is a target of dysregulation in brain endothelial cells in mice with acute respiratory SARS-CoV-2 infection, and that (2) brain endothelial cell targeted Wnt pathway activation prevents signs of cognitive impairment and neuroinflammation in acute SARS-CoV-2 infection. (3) Authors have been updated.

